## Supplemental Materials for "ResMiCo: increasing the quality of metagenome-assembled genomes with deep learning"

#### Supplementary figures

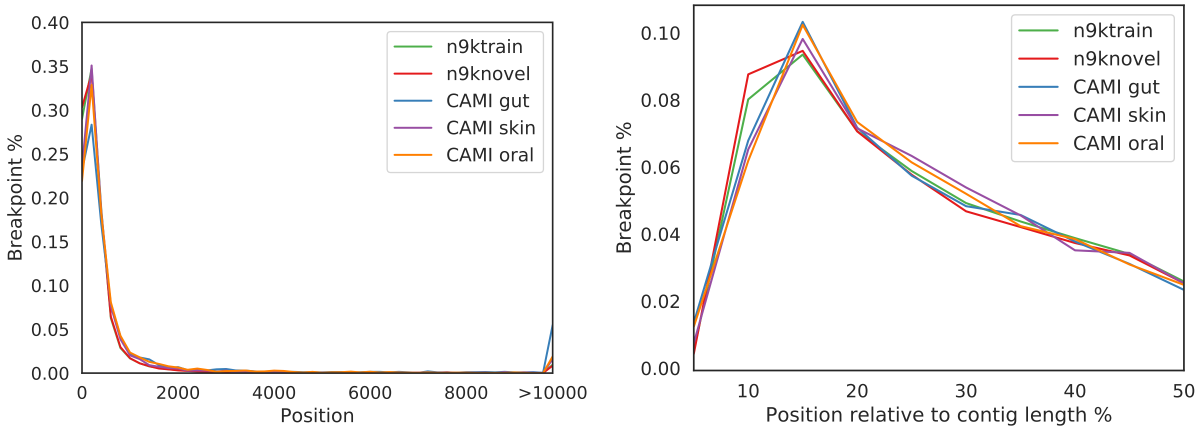


**Figure S1. Misassembly breakpoints. Left:** histogram of the absolute breakpoint location for the five datasets used for training and evaluating ResMiCo. **Right:** histogram of the breakpoint location relative to the contig length for the same datasets. Breakpoint locations are computed relative to the nearest end of the contig.

#### ResMiCo's embeddings highlight breakpoint locations

ResMiCo maintains positional information through the convolutional layers and up to the global average pooling layer, at which point the positional information is collapsed. To assess how ResMiCo's convolved features are used for classification, we visualized the output of the last residual block (Fig 2) by averaging the value of the 128 filters for each position. Formally, if we denote with $Z(\boldsymbol{x}) \in R^{128\times\lfloor|x|/8\rfloor}$ the output of the last residual block for contig $\boldsymbol{x}$, where $z_{ij}(\boldsymbol{x})$ represents the value of the $i$th filter at position $j$, we compute:

$$f_{\boldsymbol{x}}(j) = \frac{\sum_{i=1}^{128} z_{ij}(\boldsymbol{x})}{128}, j = [1..\lfloor|\boldsymbol{x}|/8\rfloor]$$

where $|\boldsymbol{x}|$ represents the length of $\boldsymbol{x}$, which is shortened by a factor of 8 due to the strided convolution at the start of the last 3 residual groups (Fig 2).

In Fig S2, we plot the resulting feature maps $f_{\boldsymbol{x}}$ for several misassembled contigs $\boldsymbol{x}$ from the *n9k-novel* dataset. In each example, the breakpoint location is associated with a spike in $f_{\boldsymbol{x}}$, with the convolutional filters spreading the signal across ≈20 embedded positions around the breakpoint, suggesting that ResMiCo can be adapted to detect misassembly locations. For misassemblies with a low ResMiCo score (false negatives), the spike is shallow (Fig S2, bottom-left) or (rarely) missing (Fig S2, bottom-right). However, we note that spikes in the feature maps are not exclusively associated with misassembled contigs. Fig S3 shows several correctly assembled contigs that feature such spikes, when ResMiCo wrongly classifies them as positives.

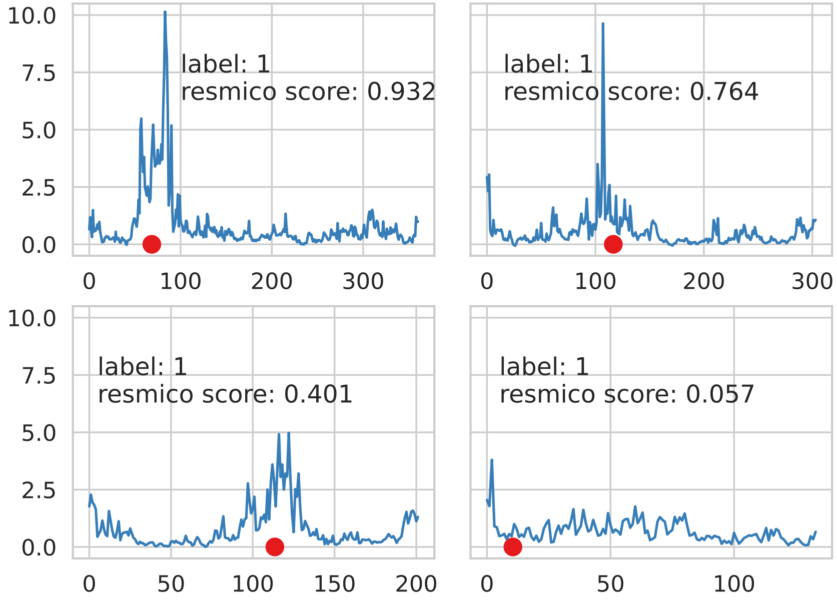


**Figure S2. Feature maps for four *n9k-novel* misassembled contigs.** The red dot marks the breakpoint location as detected by MetaQUAST. The top row shows correctly classified contigs, and the bottom row shows false negatives.

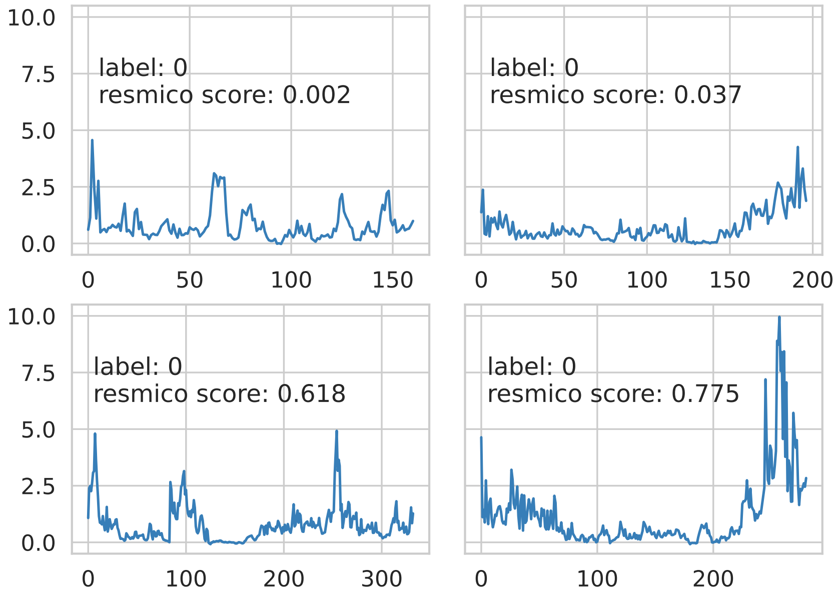


**Figure S3. Feature maps from the last layer before global pooling for four *n9k-novel* correctly assembled contigs.** The top row shows correctly classified contigs, and the bottom row shows false positives.

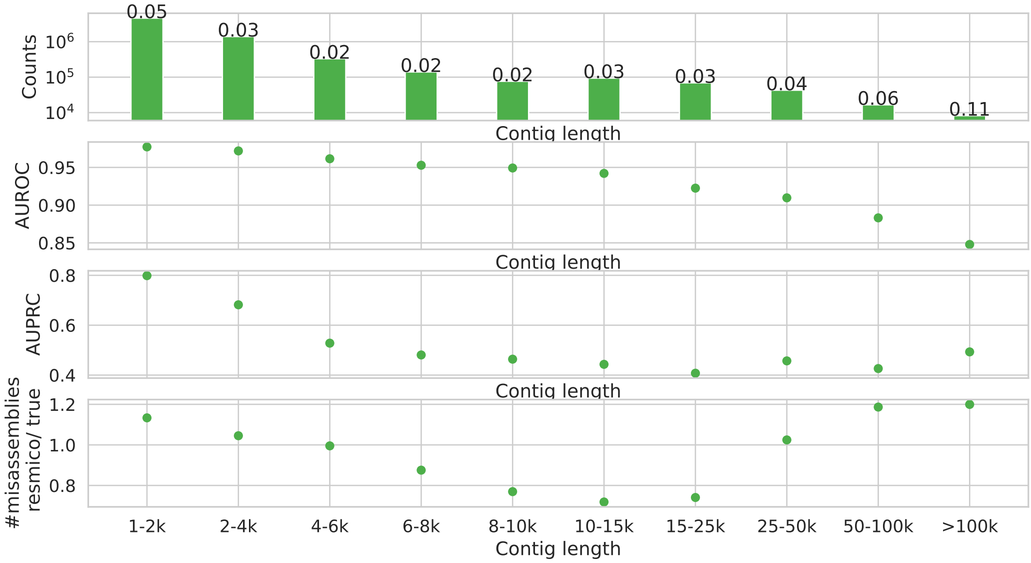


**Figure S4. The contig length distribution and ResMiCo performance on contigs of different lengths.** The contigs were grouped according to their length. On the top plot, we showed the number of contigs within a group and indicated the proportion of misassemblies with a number on top of each bar. Next, We measured AUROC and AUPRC for each group. Long contigs are more challenging than short ones. For the threshold of 0.8, we compared the number of detected misassemblies with the true number. The bottom plot shows that the ratio is close to one.

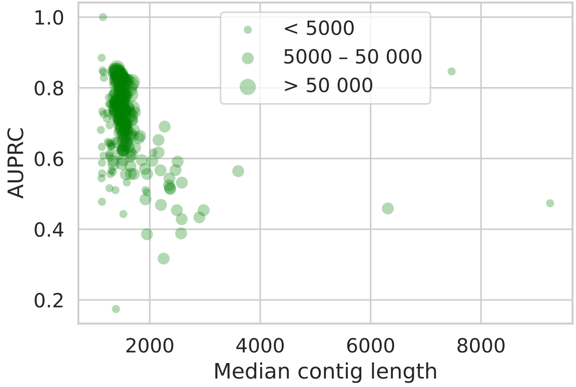


**Figure S5. AUPRC that ResMiCo achieves on 240 subsets (differing simulation parameters) of the *n9k-novel* test dataset plotted against the median length of contigs.** Size of the marker indicates a number of contigs in the subset.

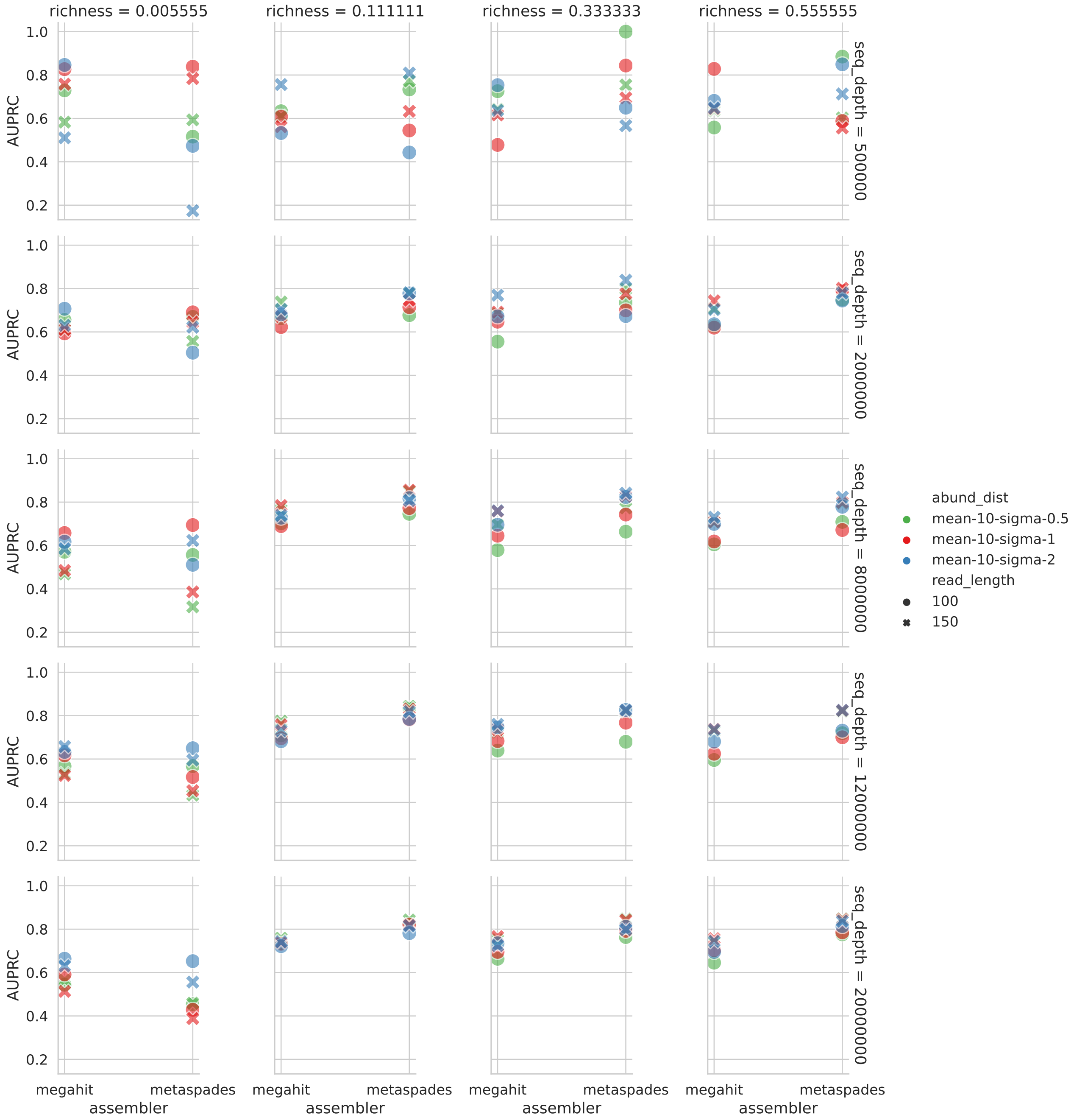


**Figure S6. ResMiCo performance measured by AUPRC on contigs from the datasets with various simulation parameters**. Simulation parameters include: community richness (fraction of 9000 genomes), genome abundance, read length, sequencing depth, and assembler. Low community richness and low sequencing depth are the most challenging conditions for ResMiCo, and the AUPRC varies substantially. For other parameter combinations, ResMiCo performance is stable, and AUPRC is between 0.6 and 0.8.

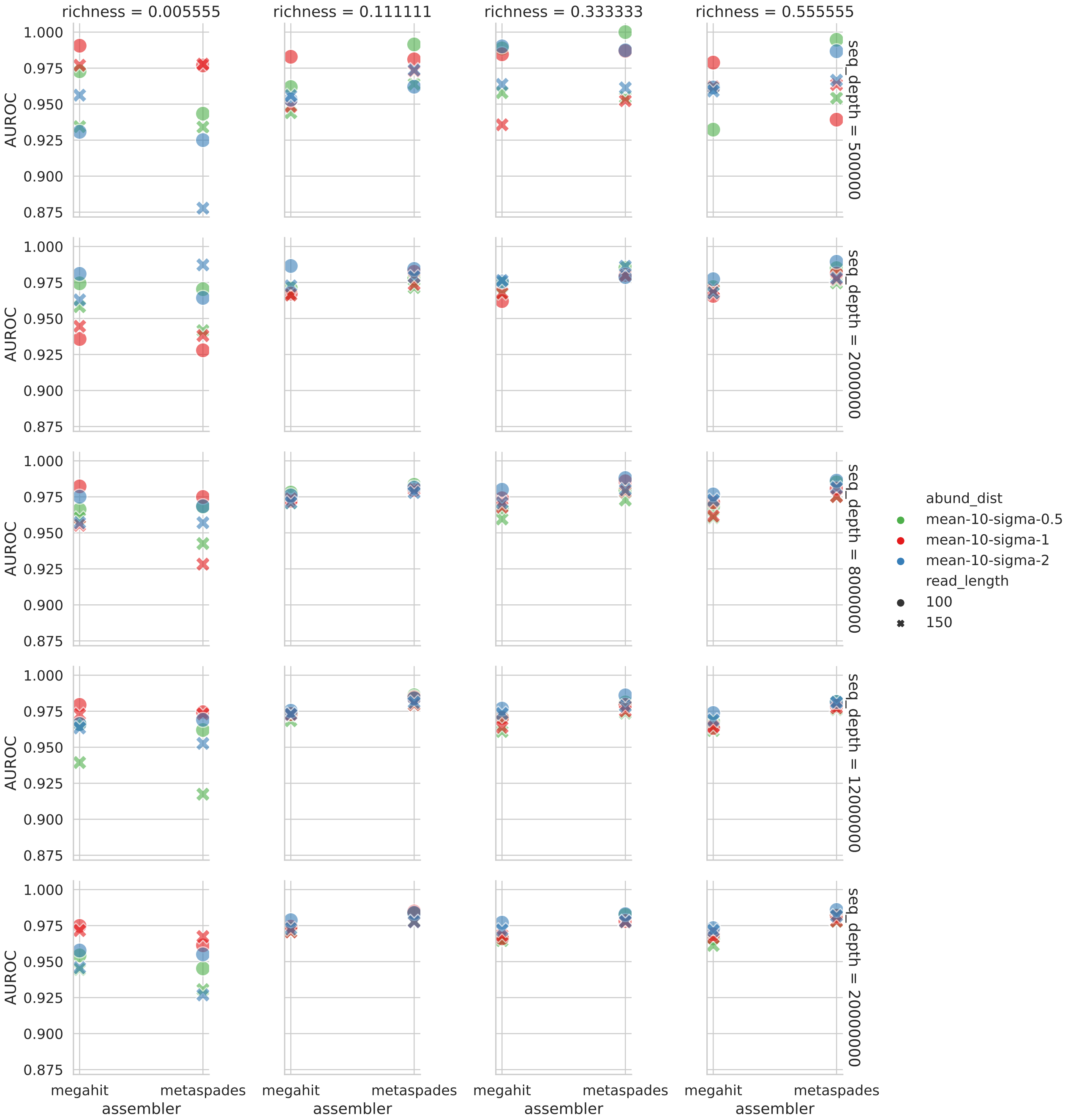


**Figure S7. ResMiCo performance measured by AUROC on contigs from the datasets with various simulation parameters** Simulation parameters include: community richness (fraction of 9000 genomes), genome abundance, read length, sequencing depth, and assembler. Low community richness and low sequencing depth are the most challenging conditions for ResMiCo. For other parameter combinations, ResMiCo performance is stable, and AUROC is around 0.97.

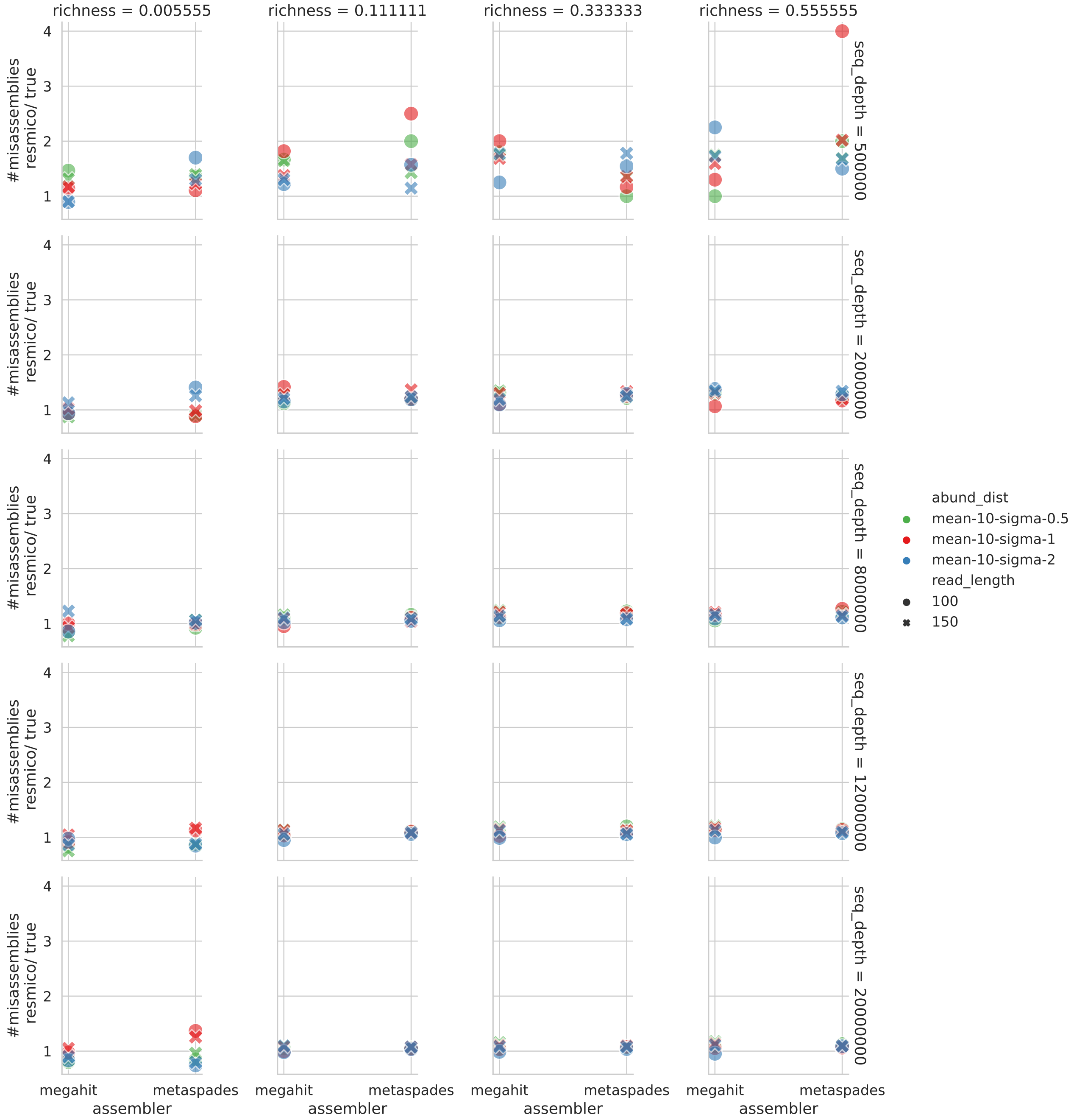


**Figure S8. Number of misassemblies found by ResMiCo divided by the true number of misassemblies in the datasets with various simulation parameters.** Simulation parameters include: community richness (fraction of 9000 genomes), genome abundance, read length, sequencing depth, and assembler. Low community richness and low sequencing depth are the most challenging conditions for ResMiCo. For other parameter combinations, ResMiCo correctly estimates the error rate using a misassembly threshold of 0.8.

### Supplementary tables

#### Features generated for ResMiCo training

| Feature | Description | Aggregation | Match | Preproc. | Used |
|---|---|---|---|---|---|
| coverage | Number of reads aligned to position | Count | A | Std | Y |
| ref_base | Nucleotide in the reference genome | - | - | Onh | N |
| num_query | Number of A/C/G/T nucleotides mapped to position | Count | A | Nrm | Y |
| num_snp | Number of aligned bases different than ref_base | Count | A | Nrm | Y |
| num_proper | Number of pair-matched aligned reads | Count | M | Nrm | Y |
| num_proper_snp | Number of SNPs from pair-matched aligned reads | Count | A | Nrm | N |
| num_discordant | Number of un-matched aligned reads | Count | A | Nrm | N |
| num_orphan | Number of orphan aligned reads | Count | M | Nrm | Y |
| al_score | Bowtie alignment scores | **Min**, **Mean**, Max, **Stdev** | M | Std | Y |
| insert_size | Length of the aligned inserts | **Min**, **Mean**, Max, Stdev | M | Std | Y |
| mapq | Bowties mapping quality of the aligned reads | Min, **Mean**, Max, Stdev | M | Std | Y |

**Table S1. The full list of positional features computed by ResMiCo pipeline.** For features computed by aggregating over multiple reads aligned to a position in a contig, the third column lists the type of aggregation, and the fourth column indicates if the feature was computed only across reads that matched the reference base (M) or for all reads (A). The fifth column states preprocessing applied to the feature: standardization (Std), normalization (Nrm), and one-hot encoding (Onh). The last column indicates if the feature was used for training the network presented in this paper: yes (Y), no (N). The corresponding type of aggregation used is in bold.

#### Model selection

The hyperparameter search for selecting the best-performing ResMiCo architecture was done by evaluating each model's performance on the validation set (i.e., randomly sampled 10% of the training set).

First, we selected the best performing neural network architecture. For this, we constructed a deep convolutional NN, a bidirectional recurrent NN with LSTM units and with GRU units, a transformer encoder, and a residual convolutional NN. All models had roughly 0.5 million trainable weights and were trained on a subset of the *n9k-train* dataset. The residual convolutional neural network outperformed the other network architectures for all contig lengths and was thus selected for this work. All subsequent hyperparameter optimizations were applied to this architecture only. We did not test all combinations of hyperparameters but rather selected the most promising ones iteratively.

#### Effect of the length cut-off and the read down-sampling on ResMiCo predictions

We studied how the down-sampling of reads and different contig length cut-offs influence contig representation and ResMiCo prediction scores. We simulated data as done for the *n9k-novel* dataset, with the following reduced set of parameters: i) richness of 1000 or 5000 genomes, ii) lognormal abundance distribution with a mean of 10 and a sigma of 1 or 2, iii) read lengths of 150bp, and iv) a sequencing depth of 2 or 8 million paired-end reads. There were a total of 96 parameter combinations. In the first experiment, we down-sampled reads to five levels (0.5, 0.1, 0.05, 0.01, 0.005) and generated features for each level. ResMiCo was then applied to the contigs shared across all levels ($n$ = 2655). Table S3 shows that the AUPRC and the ratio between the predicted error rate and the true error rate is stable across levels. Only 1% of the contigs changed prediction class across levels. In the second experiment, contigs shorter than a specified cut-off were discarded before read mapping. Contigs longer than the largest cut-off (10k) shared between all simulations were used to test ResMiCo. While 2% of contigs were assigned the opposite classes in different cut-off datasets, ResMiCo performance remained stable overall (Table S4). Taken

| Parameter | Values |
|---|---|
| Number of residual blocks (RB) in Residual groups (RG) | {[2,5,2], **[2, 5, 5, 2]**, [2, 3, 5, 5, 2], [2, 3, 5, 5, 3, 2]} |
| Number of filters for the first convolution (Conv) layer | $\{4, \mathbf{16}\}$ |
| Kernel size for Conv layers in RB | $\{3, \mathbf{5}\}$ |
| Aggregation along the spatial axis | {**global average pooling**, global max pooling, concatenated output of max and average global pooling } |
| Number of hidden units in fully connected (FC) layers $n_h$ | $\{\mathbf{50}, 100\}$ |
| Number of FC layers $n_h$ | $\{\mathbf{1}, 2\}$ |
| Initial learning rate | $\{\mathbf{0.0001}, 0.001\}$ |
| Maximum input length | $\{5000, 10000, \mathbf{20000}\}$ |

**Table S2. Hyperparameters tested for ResMiCo architecture.** The final choices for ResMiCo architecture are marked bold.

together, these results indicate that ResMiCo performance is robust to contig subsampling and filtering by contig length.

| % contigs kept | 100 | 50 | 10 | 5 | 1 | 0.5 |
|---|---|---|---|---|---|---|
| AUPRC | 0.78 | 0.78 | 0.78 | 0.78 | 0.78 | 0.78 |
| # misassemblies predicted / true | 1.21 | 1.18 | 1.22 | 1.22 | 1.21 | 1.23 |

**Table S3. Reads down-sampling effect on the ResMiCo predictions**

| Length cut-off | 1000 | 1500 | 2000 | 3000 | 5000 | 10000 |
|---|---|---|---|---|---|---|
| AUPRC | 0.38 | 0.40 | 0.39 | 0.38 | 0.38 | 0.37 |
| # misassemblies predicted / true | 0.84 | 0.80 | 0.84 | 0.80 | 0.76 | 0.80 |

**Table S4. Contigs length cut-off effect on the ResMiCo predictions**

#### ResMiCo sensitivity to changes in insert size distribution

We evaluated the robustness of ResMiCo to varying insert size distributions, given that paired-end read insert size can substantially affect assembly quality and read mapping statistics. For this experiment, we utilized the same parameter subset as done for our contig subsampling and length cutoff evaluation (see above), except we changed the insert size distributions to those shown in tables S5 and S6. We note that the *n9k-train* dataset consisted of 2 insert size distribution parameter sets: [mean=270, stdev=50] and [mean=350, stdev=75].

The AUCPR and predicted misassembly rate were quite similar to performance on the *n9k-novel* dateset, except for small insert sizes, with AUCPR dropping to 0.54 and 0.22 for mean=200 and mean=180, respectively (Table S5). Misassembly prediction rates also climbed from $< 1.18$ for mean $\geq 280$ to 2.28 and 5.39 for mean=200 and mean=180, respectively. In contrast, ResMiCo performance was quite robust to all evaluated insert size standard deviation settings (Table S6). Altogether, these results suggest that ResMiCo is sensitive to mean insert size distributions outside of the training dataset.

| Mean, Stdev of insert size distribution | 180, 50 | 200, 50 | 230, 50 | 270, 50 | 320, 50 | 380, 50 |
|---|---|---|---|---|---|---|
| AUPRC | 0.22 | 0.54 | 0.70 | 0.71 | 0.74 | 0.73 |
| # misassemblies predicted / true | 5.39 | 2.28 | 1.18 | 1.13 | 1.13 | 1.09 |

**Table S5. ResMiCo performance on the test sets with variable *mean* of the insert size distribution.** If the mean is less or equal to 200, this is an out-of-distribution sample for ResMiCo and it can not be applied to such data.

#### ResMiCo detects a 3-7% misassembly rate in real-world metagenomes

| Mean, Stdev of insert size distribution | 270, 30 | 270, 50 | 270, 70 | 270, 90 | 270, 110 |
|---|---|---|---|---|---|
| AUPRC | 0.75 | 0.71 | 0.71 | 0.72 | 0.68 |
| # misassemblies predicted / true | 1.15 | 1.13 | 1.20 | 1.22 | 1.27 |

**Table S6. ResMiCo performance on the test sets with variable *stdev* of the insert size distribution.** ResMiCo performance slightly decreases with increasing Stdev, but AUPRC remains high (0.68) up to Stdev=110.

| Dataset | Study Accession | Accession | Misassembly rate |
|---|---|---|---|
| UHGG | ERP013562 | ERS1015611 | 0.026 |
| UHGG | ERP013563 | ERS1016020 | 0.032 |
| UHGG | SRP008047 | SRS294880 | 0.034 |
| UHGG | ERP020710 | ERS1487600 | 0.035 |
| UHGG | ERP013562 | ERS1015709 | 0.037 |
| UHGG | ERP017091 | ERS1343393 | 0.037 |
| UHGG | ERP006678 | ERS537292 | 0.038 |
| UHGG | ERP010700 | ERS746721 | 0.041 |
| UHGG | ERP013562 | ERS1015695 | 0.043 |
| UHGG | ERP005860 | ERS475321 | 0.043 |
| UHGG | ERP004605 | ERS396506 | 0.043 |
| UHGG | ERP013562 | ERS1015705 | 0.045 |
| UHGG | ERP004605 | ERS396503 | 0.047 |
| UHGG | ERP014480 | ERS1076042 | 0.047 |
| UHGG | SRP093965 | SRS1820442 | 0.048 |
| UHGG | ERP012929 | ERS972211 | 0.05 |
| UHGG | SRP066479 | SRS1170732 | 0.05 |
| UHGG | ERP012929 | ERS972195 | 0.052 |
| UHGG | ERP004605 | ERS396505 | 0.056 |
| UHGG | SRP100575 | SRS1997092 | 0.06 |
| UHGG | ERP012929 | ERS953832 | 0.062 |
| UHGG | ERP005860 | ERS475197 | 0.062 |
| UHGG | SRP008047 | SRS259513 | 0.063 |
| UHGG | ERP019502 | ERS1444567 | 0.066 |
| UHGG | ERP019502 | ERS1444483 | 0.07 |
| UHGG | ERP013827 | ERS1069689 | 0.071 |
| UHGG | ERP015450 | ERS1138852 | 0.076 |
| UHGG | ERP005534 | ERS436778 | 0.077 |
| UHGG | ERP013563 | ERS1015876 | 0.08 |
| UHGG | ERP019674 | ERS1647317 | 0.081 |
| UHGG | ERP019502 | ERS1444713 | 0.09 |
| UHGG | SRP008047 | SRS259543 | 0.093 |
| UHGG | ERP017091 | ERS1343348 | 0.097 |
| UHGG | ERP006678 | ERS537234 | 0.099 |
| UHGG | ERP019674 | ERS1647292 | 0.1 |
| UHGG | ERP017091 | ERS1343328 | 0.131 |
| UHGG | ERP003612 | ERS328863 | 0.132 |
| UHGG | ERP005989 | ERS473044 | 0.132 |
| UHGG | ERP005989 | ERS473053 | 0.132 |
| UHGG | ERP019502 | ERS1444617 | 0.136 |
| UHGG | ERP005989 | ERS473285 | 0.169 |
| UHGG | ERP005989 | ERS473261 | 0.181 |
| UHGG | ERP006678 | ERS537345 | 0.2 |
| UHGG | ERP005989 | ERS473376 | 0.23 |
| TwinsUK | PRJEB40256 | ERS5053138 | 0.023 |

| | | | |
|---|---|---|---|
| TwinsUK | PRJEB40256 | ERS5053137 | 0.028 |
| TwinsUK | PRJEB40256 | ERS5053136 | 0.025 |
| TwinsUK | PRJEB40256 | ERS5053135 | 0.03 |
| TwinsUK | PRJEB40256 | ERS5053134 | 0.029 |
| TwinsUK | PRJEB40256 | ERS5053133 | 0.023 |
| TwinsUK | PRJEB40256 | ERS5053132 | 0.038 |
| TwinsUK | PRJEB40256 | ERS5053131 | 0.03 |
| TwinsUK | PRJEB40256 | ERS5053128 | 0.029 |
| TwinsUK | PRJEB40256 | ERS5053127 | 0.032 |
| TwinsUK | PRJEB40256 | ERS5053126 | 0.027 |
| TwinsUK | PRJEB40256 | ERS5053125 | 0.029 |
| TwinsUK | PRJEB40256 | ERS5053123 | 0.034 |
| TwinsUK | PRJEB40256 | ERS5053122 | 0.031 |
| TwinsUK | PRJEB40256 | ERS5053121 | 0.03 |
| TwinsUK | PRJEB40256 | ERS5053120 | 0.042 |
| TwinsUK | PRJEB40256 | ERS5053119 | 0.042 |
| TwinsUK | PRJEB40256 | ERS5053117 | 0.039 |
| TwinsUK | PRJEB40256 | ERS5053116 | 0.017 |
| TwinsUK | PRJEB40256 | ERS5053114 | 0.031 |
| TwinsUK | PRJEB40256 | ERS5053113 | 0.03 |
| TwinsUK | PRJEB40256 | ERS5053112 | 0.028 |
| TwinsUK | PRJEB40256 | ERS5053111 | 0.03 |
| TwinsUK | PRJEB40256 | ERS5053110 | 0.024 |
| TwinsUK | PRJEB40256 | ERS5053109 | 0.027 |
| TwinsUK | PRJEB40256 | ERS5053108 | 0.038 |
| TwinsUK | PRJEB40256 | ERS5053107 | 0.028 |
| TwinsUK | PRJEB40256 | ERS5053106 | 0.031 |
| TwinsUK | PRJEB40256 | ERS5053105 | 0.035 |
| TwinsUK | PRJEB40256 | ERS5053104 | 0.029 |
| TwinsUK | PRJEB40256 | ERS5053103 | 0.027 |
| TwinsUK | PRJEB40256 | ERS5053102 | 0.027 |
| TwinsUK | PRJEB40256 | ERS5053101 | 0.028 |
| TwinsUK | PRJEB40256 | ERS5053097 | 0.032 |
| TwinsUK | PRJEB40256 | ERS5053096 | 0.035 |
| TwinsUK | PRJEB40256 | ERS5053094 | 0.035 |
| TwinsUK | PRJEB40256 | ERS5053093 | 0.034 |
| TwinsUK | PRJEB40256 | ERS5053092 | 0.034 |
| TwinsUK | PRJEB40256 | ERS5053091 | 0.028 |
| TwinsUK | PRJEB40256 | ERS5053089 | 0.024 |
| TwinsUK | PRJEB40256 | ERS5053088 | 0.03 |
| TwinsUK | PRJEB40256 | ERS5053087 | 0.027 |
| TwinsUK | PRJEB40256 | ERS5053086 | 0.032 |
| TwinsUK | PRJEB40256 | ERS5053085 | 0.029 |
| TwinsUK | PRJEB40256 | ERS5053084 | 0.039 |
| TwinsUK | PRJEB40256 | ERS5053082 | 0.037 |
| TwinsUK | PRJEB40256 | ERS5053081 | 0.038 |
| TwinsUK | PRJEB40256 | ERS5053080 | 0.042 |
| TwinsUK | PRJEB40256 | ERS5053079 | 0.039 |
| TwinsUK | PRJEB40256 | ERS5053078 | 0.025 |
| TwinsUK | PRJEB40256 | ERS5053077 | 0.024 |
| TwinsUK | PRJEB40256 | ERS5053076 | 0.039 |
| TwinsUK | PRJEB40256 | ERS5053075 | 0.046 |
| TwinsUK | PRJEB40256 | ERS5053074 | 0.035 |
| TwinsUK | PRJEB40256 | ERS5053073 | 0.035 |

| | | | |
|---|---|---|---|
| TwinsUK | PRJEB40256 | ERS5053072 | 0.035 |
| TwinsUK | PRJEB40256 | ERS5053071 | 0.034 |
| TwinsUK | PRJEB40256 | ERS5053069 | 0.041 |
| TwinsUK | PRJEB40256 | ERS5053068 | 0.022 |
| TwinsUK | PRJEB40256 | ERS5053067 | 0.042 |
| TwinsUK | PRJEB40256 | ERS5053066 | 0.031 |
| TwinsUK | PRJEB40256 | ERS5053065 | 0.021 |
| TwinsUK | PRJEB40256 | ERS5053063 | 0.032 |
| TwinsUK | PRJEB40256 | ERS5053062 | 0.032 |
| TwinsUK | PRJEB40256 | ERS5053061 | 0.032 |
| TwinsUK | PRJEB40256 | ERS5053059 | 0.038 |
| TwinsUK | PRJEB40256 | ERS5053058 | 0.033 |
| TwinsUK | PRJEB40256 | ERS5053057 | 0.026 |
| TwinsUK | PRJEB40256 | ERS5053055 | 0.042 |
| TwinsUK | PRJEB40256 | ERS5053054 | 0.03 |
| TwinsUK | PRJEB40256 | ERS5053052 | 0.027 |
| TwinsUK | PRJEB40256 | ERS5053051 | 0.037 |
| TwinsUK | PRJEB40256 | ERS5053050 | 0.035 |
| TwinsUK | PRJEB40256 | ERS5053049 | 0.026 |
| TwinsUK | PRJEB40256 | ERS5053047 | 0.04 |
| TwinsUK | PRJEB40256 | ERS5053046 | 0.03 |
| TwinsUK | PRJEB40256 | ERS5053045 | 0.026 |
| TwinsUK | PRJEB40256 | ERS5053044 | 0.023 |
| TwinsUK | PRJEB40256 | ERS5053042 | 0.032 |
| TwinsUK | PRJEB40256 | ERS5053041 | 0.026 |
| TwinsUK | PRJEB40256 | ERS5053038 | 0.024 |
| TwinsUK | PRJEB40256 | ERS5053037 | 0.027 |
| TwinsUK | PRJEB40256 | ERS5053036 | 0.028 |
| TwinsUK | PRJEB40256 | ERS5053035 | 0.035 |
| TwinsUK | PRJEB40256 | ERS5053034 | 0.033 |
| TwinsUK | PRJEB40256 | ERS5053033 | 0.035 |
| TwinsUK | PRJEB40256 | ERS5053031 | 0.034 |
| TwinsUK | PRJEB40256 | ERS5053030 | 0.026 |
| TwinsUK | PRJEB40256 | ERS5053029 | 0.029 |
| TwinsUK | PRJEB40256 | ERS5053027 | 0.036 |
| TwinsUK | PRJEB40256 | ERS5053025 | 0.034 |
| TwinsUK | PRJEB40256 | ERS5053023 | 0.026 |
| TwinsUK | PRJEB40256 | ERS5053021 | 0.029 |
| TwinsUK | PRJEB40256 | ERS5053020 | 0.033 |
| TwinsUK | PRJEB40256 | ERS5053017 | 0.034 |
| TwinsUK | PRJEB40256 | ERS5053016 | 0.031 |
| TwinsUK | PRJEB40256 | ERS5053015 | 0.025 |
| TwinsUK | PRJEB40256 | ERS5053013 | 0.038 |
| Animal-gut | PRJEB38078 | ERS4537011 | 0.03 |
| Animal-gut | PRJEB38078 | ERS4537014 | 0.028 |
| Animal-gut | PRJEB38078 | ERS4537022 | 0.028 |
| Animal-gut | PRJEB38078 | ERS4537024 | 0.023 |
| Animal-gut | PRJEB38078 | ERS4537039 | 0.016 |
| Animal-gut | PRJEB38078 | ERS4537042 | 0.023 |
| Animal-gut | PRJEB38078 | ERS4537052 | 0.051 |
| Animal-gut | PRJEB38078 | ERS4537064 | 0.047 |
| Animal-gut | PRJEB38078 | ERS4537071 | 0.024 |
| Animal-gut | PRJEB38078 | ERS4537076 | 0.029 |
| Animal-gut | PRJEB38078 | ERS4537077 | 0.03 |

| | | | |
|---|---|---|---|
| Animal-gut | PRJEB38078 | ERS4537093 | 0.033 |
| Animal-gut | PRJEB38078 | ERS4537097 | 0.032 |
| Animal-gut | PRJEB38078 | ERS4537098 | 0.019 |
| Animal-gut | PRJEB38078 | ERS4537101 | 0.22 |
| Animal-gut | PRJEB38078 | ERS4537104 | 0.045 |
| Animal-gut | PRJEB38078 | ERS4537154 | 0.053 |
| Animal-gut | PRJEB38078 | ERS4537155 | 0.021 |
| Animal-gut | PRJEB38078 | ERS4537156 | 0.328 |
| Animal-gut | PRJEB38078 | ERS4537157 | 0.083 |
| Animal-gut | PRJEB38078 | ERS4537175 | 0.018 |
| Animal-gut | PRJEB38078 | ERS4537179 | 0.017 |
| Animal-gut | PRJEB38078 | ERS4537189 | 0.053 |
| Animal-gut | PRJEB38078 | ERS4537203 | 0.026 |
| Animal-gut | PRJEB38078 | ERS4537227 | 0.059 |
| Animal-gut | PRJEB38078 | ERS4537248 | 0.032 |
| Animal-gut | PRJEB38078 | ERS4537271 | 0.018 |
| Animal-gut | PRJEB38078 | ERS4537272 | 0.016 |
| Animal-gut | PRJEB38078 | ERS4537273 | 0.052 |
| Animal-gut | PRJEB38078 | ERS4537280 | 0.024 |

**Table S7. ResMiCo estimates the prevalence of misassembled contigs in publicly available metagenomic data.**

### Supplementary Methods

#### Optimizing data generation performance

ResMiCo's data simulation pipeline generates features for training the ResMiCo model by re-aligning reads against the putative contigs and extracting statistics for each positions from the realigned reads (Fig 1, d). Since the original implementation of the feature generation pipeline, based on pysam+htslib, was prohibitively slow for manipulating the amount of data that ResMiCo required for training, we wrote our own pileup generation and feature extraction library. ResMiCo's feature generation is based on the BamTools (`https://github.com/pezmaster31/bamtools`) library that provides raw access to the BAM files generated by Bowtie2 [25]. The speed gain was achieved by loading all alignments in memory into re-sizable chunks (based on the machine's available RAM) rather than streaming data from disk. Also, rather than first generating a pileup and then analyzing the resulting pileup file, the two steps were unified (i.e., the features were computed on the fly at the time of the pileup generation). The contigs in BAM files were read and processed in parallel (since they are independent of each other), thus making effective use of all CPU cores. Writing data to disk was delegated to a separate consumer thread, thus avoiding slow IO blocking pileup generation. In addition, the resulting features were written on disk in a binary format with a fixed schema, which allowed both efficient storage and indexing. The modified feature generation was about an order of magnitude faster than the original pysam+htslib-based feature generation. The source code for the feature generation library is available at `https://github.com/leylabmpi/ResMiCo/tree/master/ResMiCo-SM/feature_extractor`.
